# A dual MTOR/NAD+ acting gerotherapy

**DOI:** 10.1101/2023.01.16.523975

**Authors:** Jinmei Li, Sandeep Kumar, Kirill Miachin, Nicholas L. Bean, Ornella Halawi, Scott Lee, JiWoong Park, Tanya H. Pierre, Jin-Hui Hor, Shi-Yan Ng, Kelvin J. Wallace, Niklas Rindtorff, Timothy M. Miller, Michael L. Niehoff, Susan A. Farr, Rolf F. Kletzien, Jerry Colca, Steven P. Tanis, Yana Chen, Kristine Griffett, Kyle S. McCommis, Brian N. Finck, Tim R. Peterson

## Abstract

The geroscience hypothesis states that a therapy that prevents the underlying aging process should prevent multiple aging related diseases. The mTOR (mechanistic target of rapamycin)/insulin and NAD+ (nicotinamide adenine dinucleotide) pathways are two of the most validated aging pathways. Yet, it’s largely unclear how they might talk to each other in aging. In genome-wide CRISPRa screening with a novel class of N-O-Methyl-propanamide-containing compounds we named BIOIO-1001, we identified lipid metabolism centering on SIRT3 as a point of intersection of the mTOR/insulin and NAD+ pathways. In vivo testing indicated that BIOIO-1001 reduced high fat, high sugar diet-induced metabolic derangements, inflammation, and fibrosis, each being characteristic of non-alcoholic steatohepatitis (NASH). An unbiased screen of patient datasets suggested a potential link between the anti-inflammatory and anti-fibrotic effects of BIOIO-1001 in NASH models to those in amyotrophic lateral sclerosis (ALS). Directed experiments subsequently determined that BIOIO-1001 was protective in both sporadic and familial ALS models. Both NASH and ALS have no treatments and suffer from a lack of convenient biomarkers to monitor therapeutic efficacy. A potential strength in considering BIOIO-1001 as a therapy is that the blood biomarker that it modulates, namely plasma triglycerides, can be conveniently used to screen patients for responders. More conceptually, to our knowledge BIOIO-1001 is a first therapy that fits the geroscience hypothesis by acting on multiple core aging pathways and that can alleviate multiple conditions after they have set in.

**Brief Summary:** These studies characterize a novel gerotherapy, BIOIO-1001, that identifies lipid metabolism as an intersection of the mTOR and NAD+ pathways.

## INTRODUCTION

Phenotypic screening, sometimes serendipitously performed, has been how many of the most influential drugs were identified ^1^. Interestingly, phenotypic screening fell out of favor as the molecular biology revolution and rational drug design – developing drugs based on knowledge of a target – gained hold. A major problem with phenotyping screening for those in drug development has been that typically one does not learn their mechanism of action (MoA) from them. Fortunately, phenotyping screening has made a comeback as new methodologies such as CRISPR have made drug target deconvolution easier ^2^.

Another issue rational drug design proponents have had is that compounds that act on multiple targets are considered sub-optimal for drug development. The one target-one disease creates a simple narrative that scientists, publishers, funders, and regulators have been able to get behind. Yet, these traditional pharma forces are at odds with the aging field both in that: 1) a strongly performing gerotherapy would likely have more pervasive effects than a highly selective single target drug could produce ^3^ and 2) aging is not a FDA approved disease indication. It is an exhausting challenge to gain drug approval for one indication, which has made it prohibitive for researchers to know whether their drug might work in additional contexts.

Taking an unbiased genome-wide screening approach led us to the determination that a novel compound series we identified, BIOIO-1001, is a multi-pathway, multi-disease therapy. Molecularly speaking, because BIOIO-1001 acts at the intersection of two canonical aging pathways, mTORC1 (mechanistic target of rapamycin) and NAD+ metabolism, both which act pervasively on the hallmarks of aging ^3, 4^, our work suggests BIOIO-1001 is a bona fide novel gerotherapy.

## RESULTS

### Identification of SIRT3 as a BIOIO-1001 genetic target

With the premise that insulin resistance is important to many diseases of aging ^5^, phenotypic screening was performed to identify novel insulin sensitizing agents. From these screens, a compound series was identified that we named BIOIO-1001 (**Fig. 1A**). The BIOIO-1001 series are N-O-Methyl-propanamide-containing compounds derived from the thiazolidinedione (TZD) ^6^, pioglitazone, that lack the heterocyclic C_3_NS ring that defines the TZD compounds. In vitro testing established that BIOIO-1001 did not activate the canonical TZD target, PPARgamma (**Fig. S1A**) ^7^. BIOIO-1001 also did not inhibit the mitochondrial pyruvate carrier or activate a panel of nuclear hormone receptors (**Fig. S1B, C**) – factors which might have been involved in the MoA of BIOIO-1001 based on our and others previous work ^8^.

**Figure 1.**
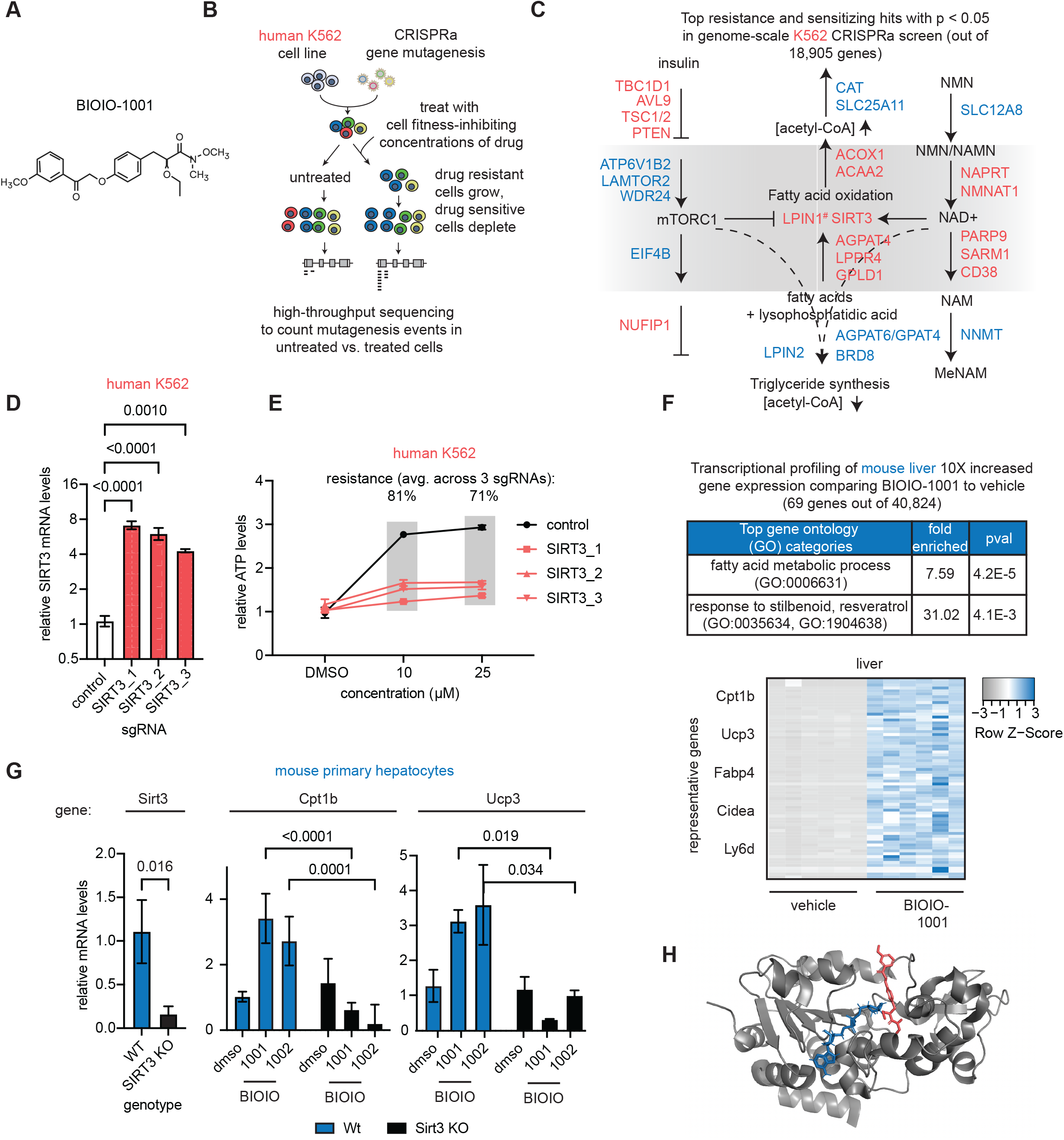
Genome-wide screening with BIOIO-1001 identifies SIRT3 as a point of intersection between the MTOR and NAD+ pathways. **A**. Chemical structure of BIOIO-1001. **B**. Schematic of a CRISPRa screen with BIOIO-1001. Screening was performed in duplicate with 50µM of BIOIO-1001 or vehicle (DMSO) in K562 cells. **C**. Genetic map of top hits from the BIOIO-1001 CRISPRa screen. Sensitizing and resistance hits (p < 0.05, based on the average phenotype of top three sgRNAs out of 10) are shown in red and blue, respectively. The putative mechanism of action of BIOIO-1001 is where the hits flip from resistance to sensitizing to resistance or vice versa (highlighted in gray) in both the mTORC1 and NAD+ pathways. The common target of both pathways is fatty acid oxidation, and specifically the mTORC1 and NAD+ targets, LPIN1 and SIRT3, respectively. Asterisk on LPIN1 indicates it was sensitizing but its p value was greater than 0.05. **D**. Relative SIRT3 mRNA expression in SIRT3 CRISPRa K562 cells. SIRT3 mRNA levels were assessed by RT-qPCR. One-way ANOVA for all SIRT3 sgRNAs vs. control sgRNA. **E**. Relative ATP levels in SIRT3 overexpressing and control cells in the presence of the indicated concentrations of BIOIO-1001 or vehicle (DMSO). ATP levels were measured after 72 hours treatment in cells from D. One-way ANOVA for all SIRT3 sgRNAs vs. control sgRNA. **F**. Transcriptionally profiling of livers harvested from mice treated with BIOIO-1001. Mice fed a high fat diet for 12 weeks were gavaged once daily with 30mg/kg/day of BIOIO-1001 or vehicle (n=6) for one week prior to euthanasia. Genes that were increased in expression 10-fold compared to vehicle treated condition were included in the gene ontology analysis. Only genes with counts per million (CPM) in the vehicle treated condition of greater than 0.1 were included. **G**. Wildtype and SIRT3 deficient primary mouse hepatocytes treated with BIOIO-1001, BIOIO-1002, or vehicle at the indicated concentrations for 24 hours. Gene expression of the indicated genes were measured. One-way ANOVA for all SIRT3 sgRNAs vs. control sgRNA. Asterisk (*) indicates p < 0.05 (n=4). **H**. Predicted small molecule docking pose of BIOIO-1001 with SIRT3. Shown is the highest confidence docking pose of BIOIO-1001 (red) with human SIRT3 protein (PDB: #5514, solvent and PEG removed). NAD-Ribose (blue) is visible bound in the active site of SIRT3. Zinc ion (gray) shown.

We and others have previously used genome-scale CRISPR-based screening in pinpointing a small molecule’s mechanistic target based on its pattern of resistance and sensitizing hits ^9-11^. To deconvolute the MoA of BIOIO-1001, we performed a genome-scale CRISPR activation (CRISPRa) screen with BIOIO-1001 (**Fig. 1B**). This screen pinpointed downregulation of the mTORC1/insulin and upregulation of the NAD+ pathways, respectively, as most important to BIOIO-1001’s action (**Fig. 1C, Supplemental Table 1**). This is consistent with the sign convention of effects of these two pathways in promoting longevity. The hits flipped from resistance to sensitizing to resistance or vice versa for each pathway in a way that centered on BIOIO-1001 promoting fatty acid oxidation (FAO) (Fig. 1C). Both mTORC1 and NAD+ have effectors that contribute significantly to fatty acid oxidation, namely LPIN1 and SIRT3 respectively ^12-15^. LPIN1 was directionally consistent in the CRISPRa screen with its positive role in FAO ^12^, but it was not a statistically significant hit. Whereas SIRT3 scored strongly and follow up analysis validated it as a BIOIO-1001 genetic target (**Fig. 1D, E**). Notably, CRISPRa mediated increased SIRT3 mRNA expression led to increased LPIN1 mRNA expression, which is consistent with these two genes acting in concert (**Fig. S1D)**. This has analogies to the results for LPIN2 in the CRISPRa screen. LPIN2 suppresses LPIN1 expression and indeed LPIN2 was a strong hit, but in the opposite direction of LPIN1 (Fig. 1C) ^16^. Phosphatidate phosphatase activity (GO:0008195) was a top gene ontology category in transcriptional profiling of BIOIO-1001 treated K562 cells based on using Enrichr ^17^, which identifies pathway, disease, and drug signatures that intersect with an input gene list (**Fig. S1E, Supplemental Table 1**). The identification of fatty acid oxidation-related disease biology (short chain acyl-CoA dehydrogenase deficiency, ORPHA:26792) as a top gene signature in unbiased genome wide transcriptional profiling corroborated the similar finding in our unbiased genome wide CRISPRa screen (Fig. S1E, Supplemental Table 1). More evidence that LPIN1 appears to be involved in the BIOIO-1001 MoA is that BIOIO-1001 increased the levels of the LPIN1 substrate, phosphatidic acid, whereas it decreased the levels of the LPIN1 product, diacylglycerol, in livers of mice treated with the drug (**Fig. S1F**).

Since K562 is a cancer cell line we sought a more physiological relevant context to characterize BIOIO-1001’s molecular mechanisms. Transcriptional profiling of mice treated with BIOIO-1001 revealed fatty acid metabolic process (GO:0006631) as the top gene signature, consistent with our results in K562 cells (**Fig. 1F, Supplemental Table 1**). Validation of ex vivo mouse liver confirmed SIRT3 not only as a strong genetic interactor with BIOIO-1001, but more importantly as required for metabolic signaling downstream of both mTOR and NAD+ (**Fig. 1G**) ^14, 15^. The BIOIO-1001 back up compound, BIOIO-1002, also required SIRT3 for its metabolic effects (specifically, to activate the expression of Cpt1b and Ucp3 in mouse hepatocytes, Fig. 1G). Interestingly, the top signature in our liver transcriptional profiling of high-fat diet fed mice based on fold change was responses to stilbenoid and resveratrol (GO:0035634, GO:1904638). Resveratrol is a well-known stilbenoid that has long been connected to NAD+ metabolism and specifically, the sirtuins ^18^. Though the mechanism(s) by which resveratrol acts on the sirtuins are controversial, it has been shown that resveratrol can bind directly to SIRT1 ^19^. These results led us to hypothesize that potentially BIOIO-1001 might bind SIRT3. In silico docking of BIOIO-1001 to SIRT3 predicted a singular interaction between BIOIO-1001 to SIRT3 outside of the NAD-Ribose binding pocket (**Fig. 1H, Fig. S1G**). Taken together, the above suggests the BIOIO-1001 series is a novel metabolic regulator of lipid metabolism via its mTOR and NAD+ targets, LPIN1 and SIRT3.

### BIOIO-1001 provides in vivo and in vitro protection against high fat diet and NASH models

The mTOR and NAD+ pathways have broad metabolic effects ^20, 21^. To test these effects in vivo, we put mice on a high fat diet (HFD) for 10 weeks and then treated mice with either BIOIO-1001 or the aforementioned insulin sensitizer benchmark, pioglitazone (PIO), for 10 days (**Fig. S2A**). The rationale for this protocol rather than co-administering the diet and drug at the start was to model a real-world scenario where the therapy would be given after disease has set in. Both BIOIO-1001 and PIO improved glucose tolerance in HFD fed mice (**Fig. S2B**). Feeding mice HF diet increased plasma insulin concentration, but consistent with an insulin sensitizing effect, insulin concentration in HF-fed mice treated with BIOIO-1001 or PIO were significantly lower than HF vehicle controls and not different than LF diet-fed controls (**Fig. S2C**). This is likely due to reduced insulin secretion, rather than clearance, because C-peptide concentrations in the blood were also reduced by treating with BIOIO-1001 or PIO (Fig. S2C). This suggested that less insulin was needed to maintain normoglycemia. Plasma non-esterified fatty acids and liver triglycerides were both reduced by BIOIO-1001 or PIO (**Fig. S2C, D**), but interestingly, plasma triglycerides were uniquely reduced by BIOIO-1001, but not by PIO (Fig. S2C). A subset of mice from each group was injected with saline or insulin prior to sacrifice and muscle insulin signaling was assessed. BIOIO-1001 and PIO increased the phosphorylation of AKT in response to insulin stimulation (**Fig. S2E**). These data suggest that BIOIO-1001 improves metabolic abnormalities and insulin sensitivity in obese mice.

Inclusion of fructose and cholesterol in high fat diets can exacerbate diabetes and fatty liver and lead to non-alcoholic steatohepatitis (NASH), which is characterized by inflammation and fibrosis ^22, 23^. We put mice on a diet high in trans-fat, fructose, and cholesterol (HTF-C) as a mouse model of NASH. Like the HF diet, we provided the HTF-C diet for a prolonged period (16 weeks) before briefly treating the mice with BIOIO-1001 or PIO (last three weeks) (**Fig. 2A**). Strikingly, despite BIOIO-1001 being given for < 20% of the time the mice were on the HTF-C diet, it significantly reduced liver injury and a host of NASH-related phenotypes, including (plasma transaminases - ALT, AST) and liver triglyceride content more strongly than PIO (**Fig. 2B-D**). BIOIO-1001 also tended to improve NAS and fibrosis scoring after this short treatment period (**Fig. 2E-F**). Consistent with these histologic and plasma findings, BIOIO-1001 was more protective against inflammation and fibrosis markers at the transcriptional level than PIO (**Fig. 2G-I**). Ex vivo BIOIO-1001 reduced several inflammation and fibrosis markers in a SIRT3 dependent manner (**Fig. 2J**). Taken together, these results suggest BIOIO-1001 alleviates numerous diet-induced phenotypes found in NASH in a SIRT3 dependent manner.

**Figure 2.**
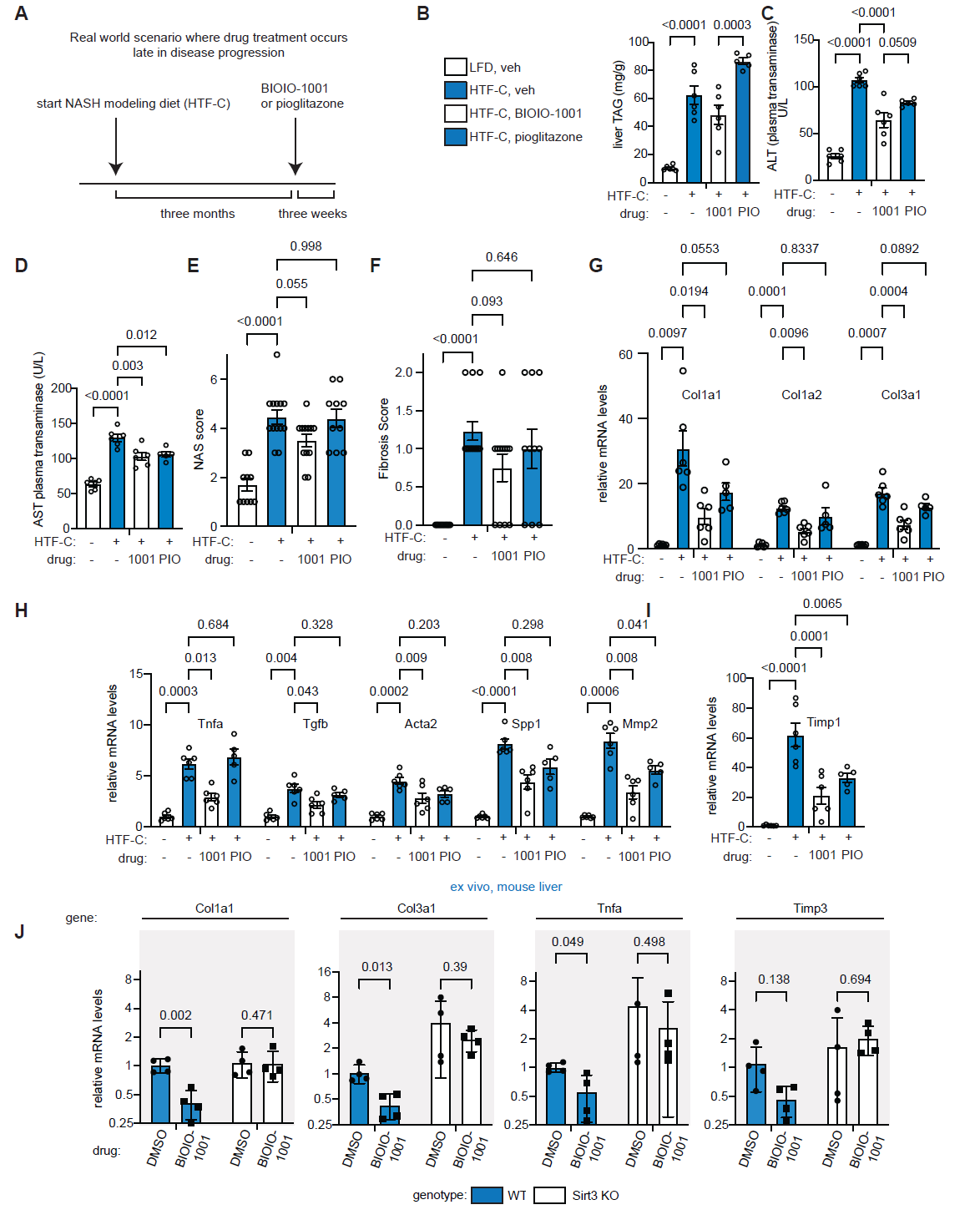
BIOIO-1001 reverses liver injury and stellate cell activation in a mouse model of NASH. **A**. Schematic depicting the time course of diet and drug treatment. Mice were fed a low fat or HTF-C diet for 16 weeks and then treated with vehicle (veh), PIO, or BIOIO-1001 for 3 weeks by gavage. **B-F**. Liver triglycerides (B), circulating ALT activity (C), circulating AST (D), hepatic NAS (E) and fibrosis scoring (F) from mice treated in A. One-way ANOVA was performed to compare the four groups (n=6 except, NAS and fibrosis where n=10-13). **G-I**. Expression of genes encoding markers of inflammation and stellate cell activation in mice treated as in A. **J**. Wildtype and SIRT3 deficient primary mouse hepatocytes treated with BIOIO-1001 or vehicle at the indicated concentrations for 24 hours. Gene expression of the indicated genes were measured. P-values based on multiple unpaired t-tests.

### BIOIO-1001 provides in vivo and in vitro protection against sporadic and familial models of ALS

We performed transcriptional profiling on several tissues (heart, liver, adipose tissue) from the mice from our HF diet study. These experiments identified long chain fatty acid metabolism as a core signature across all three tissues as increased by BIOIO-1001 (**Supplemental Table 2**). Long chain fatty acid metabolism is selectively deregulated in SIRT3 deficient mice ^13^. Interestingly, based on the dbGaP human database of genotypes and phenotypes ^24^ our gene list most strongly was comprised of genes (e.g., RNF144A, RD3, KCNMA1, SUSD1, p = 0.0014) involved in amyotrophic lateral sclerosis (ALS) (**Fig. 3A**). This was interesting because these genes are involved in inflammation and fibrosis ^25-28^ and inflammation and fibrosis are key consequences of altered metabolism in both NASH and ALS pathology ^23, 29, 30^.

**Figure 3.**
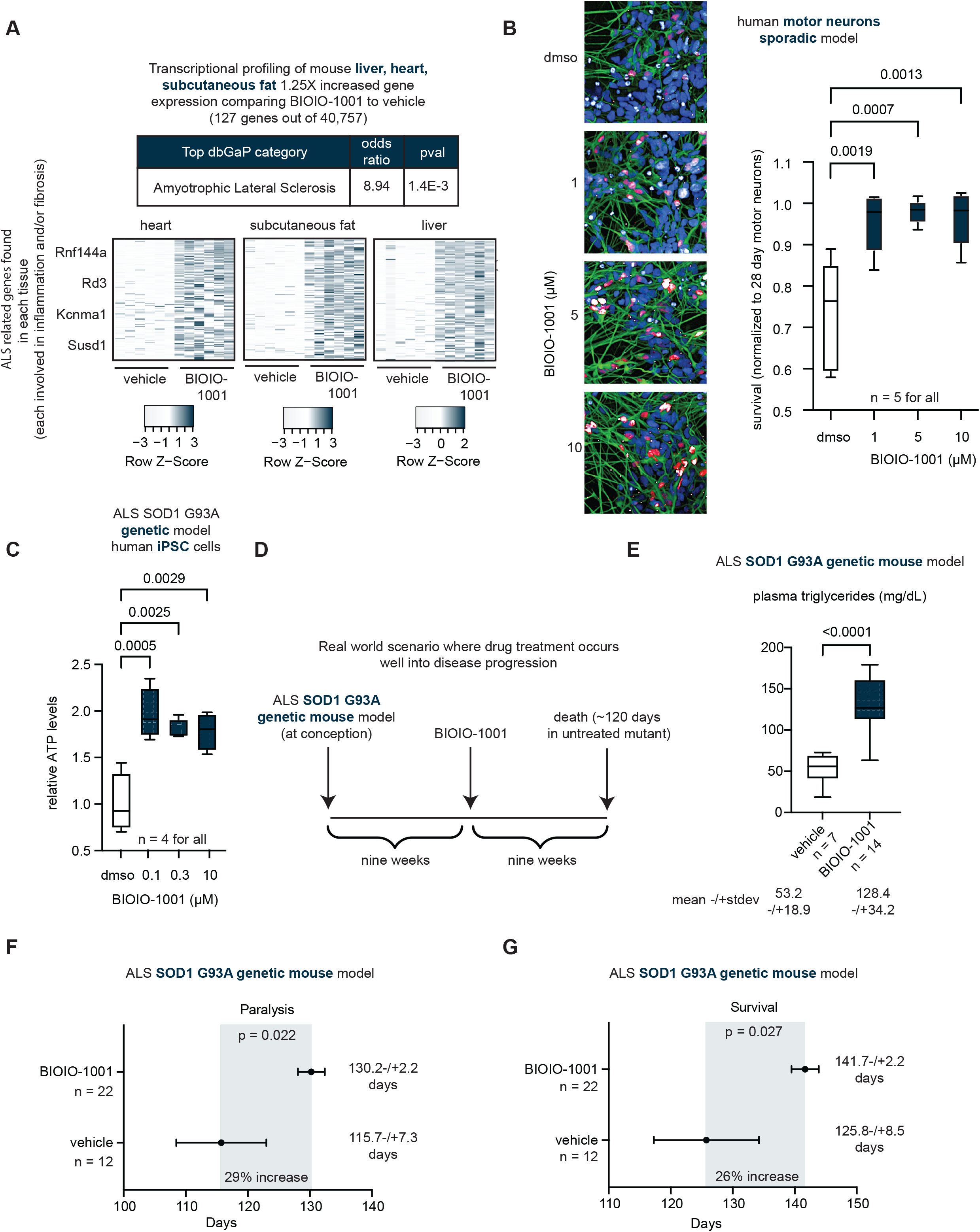
BIOIO-1001 promotes cell and organismal survival in genetic and sporadic models of ALS. **A**. Transcriptional profiling (RNAseq) in liver, heart, and subcutaneous fat using BIOIO-1001. Top gene signatures from dbGaP via Enrichr were derived from those genes that were 1.25X increased due to 30mg/kg BIOIO-1001 as in Figure 2F (127 genes out of 40,757). **B**. The indicated concentrations of BIOIO-1001 or DMSO were given to iPSCs derived from a patient with sporadic ALS. 31-day motor neuron survival as judged by ISL1 (motor neuron marker, in red)/DAPI (nuclear marker, in blue) ratios were normalized to day 28 motor neuron survival. SMI-32, another neuronal marker is in green. **C**. Cell viability in human iPSCs containing a G93A SOD1 mutation. Cells were treated with the indicated concentrations of BIOIO-1001 or vehicle for 24hrs. **D**. Schematic depicting the time course of BIOIO-1001 treatment in the hereditary (genetic) ALS mouse model, G93A Sod1. **E**. Plasma triglycerides collected from animals treated as in D after four weeks of treatment. **F, G**. Paralysis and survival analysis of G93A Sod1 mutant mice treated with BIOIO-1001 or vehicle as schematized in D.

We investigated further a potential role for BIOIO-1001 in treating ALS. There are two forms of ALS, sporadic and familial, with sporadic accounting for 90% of all cases ^31^. In ALS, motor neuron survival is impaired, and this contributes to paralysis and ultimately the death of the patient. We measured the effects of BIOIO-1001 on the survival of motor neurons derived from sporadic ALS patients. In these assays BIOIO-1001 was protective towards cell viability (**Fig. 3B**).

To study hereditary ALS, we performed analogous experiments as in Fig. 3B using cells with the well-known SOD1-G93A mutation ^32^. The results were similar to those in Fig. 3B – BIOIO-1001 was protective (**Fig. 3C**). We tested BIOIO-1001 in vivo in the SOD1-G93A mouse model of the familial ALS condition ^32^. This commonly used model displays many features of ALS including neuromuscular degeneration leading to paralysis and shortened lifespan. We deployed the same paradigm as with the NASH model. That is, we treated mice with BIOIO-1001 after disease set at nine weeks (typical lifespan is 18 weeks with this model) to mimic the real-world scenario that the drug would be given to symptomatic patients (**Fig. 3D**). Interestingly, compared with vehicle treatment, BIOIO-1001 increased plasma triglycerides in the SOD1-G93A mice (**Fig. 3E**). Unlike in metabolic diseases like NASH, in ALS, higher triglycerides levels have been shown to correlate with better prognosis in humans ^33^. Indeed, BIOIO-1001 was protective against paralysis and prolonged the lifespans of the SOD1-G93A mice by 29% and 26%, respectively (**Fig. 3F, G**). Taken together with our in vitro results, these results suggest BIOIO-1001 alleviates numerous phenotypes found in ALS.

## DISCUSSION

The geroscience hypothesis states that a therapy that checks the underlying aging process should prevent multiple aging related diseases ^34^. Herein we provide evidence BIOIO-1001 is a novel gerotherapy that modulates multiple aging pathways and can be therapeutic in multiple age-related diseases.

There are several core pathways that affect aging. Rapamycin targets the mTOR pathway. NAD+ metabolites boost the NAD+ pathway. Our work presents insights into how the mTOR and NAD+ might talk to each other in aging. Specifically, we identify lipid metabolism and the genes, LPIN1 and SIRT3, as the point of intersection of the mTOR and NAD+ pathways. That BIOIO-1001 enabled this discovery speaks to the value of using drugs as a tool to understand biology. That BIOIO-1001 enabled this discovery speaks to the value of using drugs in making biological discovery. Mechanistically, we speculate that BIOIO-1001 exerts its action epigenetically by affecting the levels of acetyl-CoA (Fig. 1C). BIOIO-1001 is a N-O-Methyl-propanamide compound that is hydroxamic-like. Hydroxamic compounds are known to have epigenetic effects ^35^. Also, both LPINs and SIRTs affect acetylation of lipid metabolism genes ^36, 37^. In future studies, it will be interesting to determine the precise mechanism of BIOIO-1001’s activity on the LPINs and SIRTs.

Both inhibition of mTORC1 and activation of NAD+ effectors suppress inflammation and fibrosis ^38, 39^. In future studies it will be interesting to dissect the mechanisms by which the effect of BIOIO-1001 on lipid metabolism translate to reduced inflammation and fibrosis. Additionally, it will be interesting to determine how decreased vs. increased plasma triglycerides can lead to improved outcomes in NASH vs. ALS models as we observed.

There are several issues with the animal models used in aging research. Lifespan studies do not inform how a therapy might work in disease. Progeria models are too extreme in how short lived they are and too few people have this condition for it to be considered relevant to more common aging-related diseases ^40^. Thus, we propose ALS could be an interesting new gerotherapy proving ground. Like aging itself, ALS has many unrelated molecular targets. If a therapy works in diverse ALS models, we hypothesize it could work across diseases. Our lead asset, BIOIO-1001, provides proof of concept of this hypothesis.

## Supporting information

BIOIO-1001-SuppTable1

BIOIO-1001-SuppTable2

## Abbreviations

SIRT3: Sirtuin 3
LPIN1: Lipin 1
MTOR: mechanistic target of rapamycin
NAD+: Nicotinamide adenine dinucleotide
NASH: non-alcoholic steatohepatitis
ALS: amyotrophic lateral sclerosis
SOD1: super oxide dismutase

---

**Figure S1.**
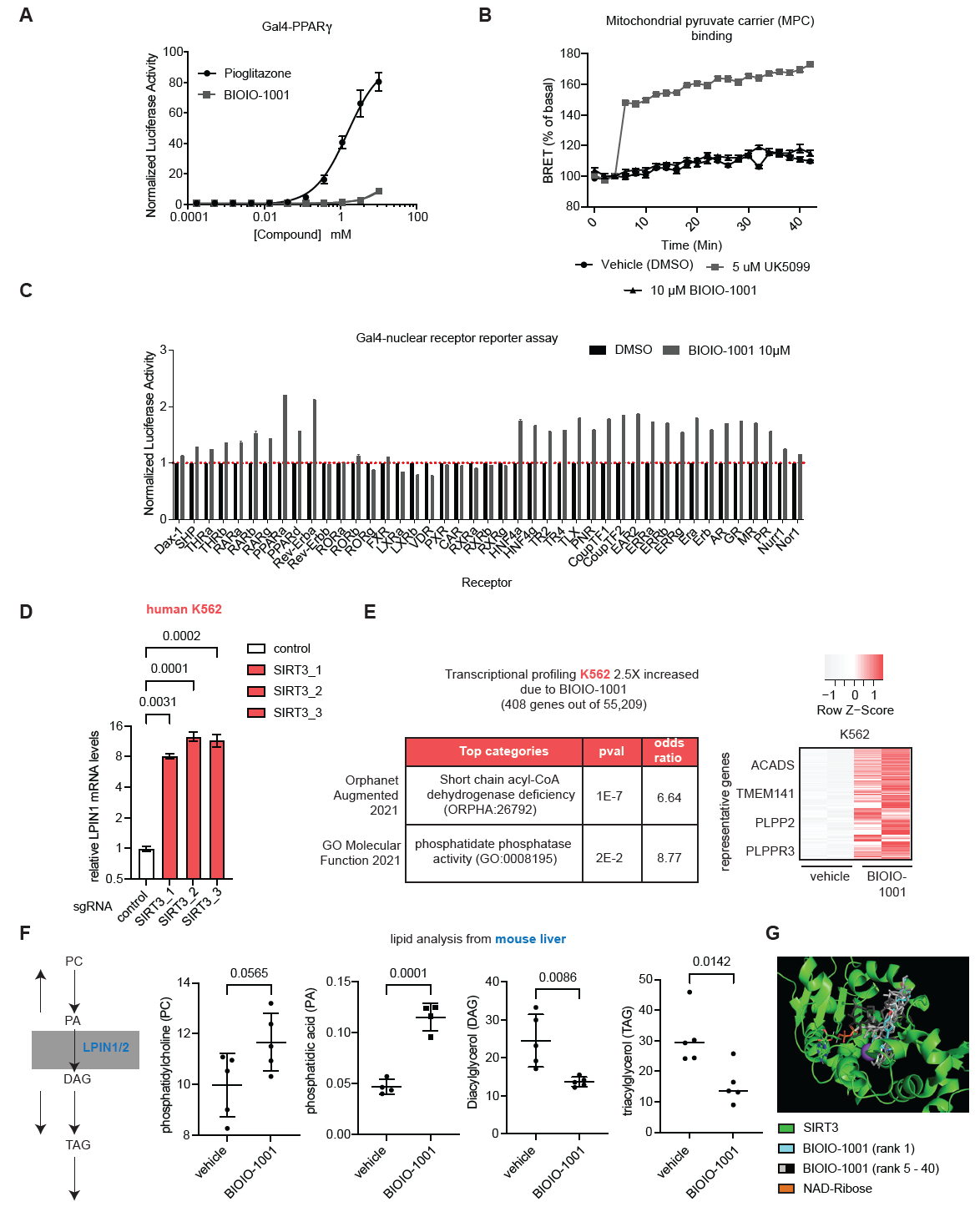
Data related to Figure 1. **A**. PPAR gamma (*γ*) reporter assay in the presence of the indicated concentrations BIOIO-1001 or pioglitazone. **B**. Effect of BIOIO-1001 and BIOIO-1002 on a BRET (bioluminescence resonance energy transfer)-based assay for inhibitors of the mitochondrial pyruvate carrier (RESPYR). After establishing basal activity values, BIOIO-1001, BIOIO-1002, or the MPC inhibitor UK5099 were added to cells expressing the RESPYR proteins at the four-minute time point. Values are presented as mean ± standard error of the mean. *p<0.05 UK5099 compared to DMSO at all timepoints. **C**. The graph depicts Gal4-responsive luciferase activity in cells expressing fusion proteins of Gal4 and listed nuclear receptors. Cells expressing these proteins were treated with vehicle or BIOIO-1001 at 10*μ*M. **D**. Relative LPIN1 mRNA expression in SIRT3 CRISPRa K562 cells. LPIN1 mRNA levels were assessed by RT-qPCR. One-way ANOVA for all SIRT3 sgRNAs vs. control sgRNA. **E**. Transcriptional profiling (RNAseq) in K562 cells using BIOIO-1001. Top gene signatures from Enrichr were derived from those genes that were 2.5X increased due to 50µM BIOIO-1001 (408 genes out of 55,209). **F**. Analysis of lipids regulated by phosphatidic acid phosphatases from mouse livers. Mice were fed high fat (60% fat) diet for 10 weeks and then gavaged daily with 30mg/kg/day of BIOIO-1001 or vehicle for one week prior to euthanasia. Total phosphatidylcholine, phosphatidic acid, diacylglycerol, and triacylglycerol were extracted and analyzed by mass spectrometry. Schematic depicting Lipins in the triglyceride synthesis pathway is shown. **G**. Set of predicted small molecule docking poses of BIOIO-1001 with SIRT3. Shown are the highest confidence-score docking pose of BIOIO-1001 (cyan) as well as the lower rank predictions (grey to black) with human SIRT3 protein (PDB: #5514, solvent and PEG removed). NAD-Ribose (orange/green) is visible bound in the active site of SIRT3. Zinc ion (purple) shown.

**Figure S2.**
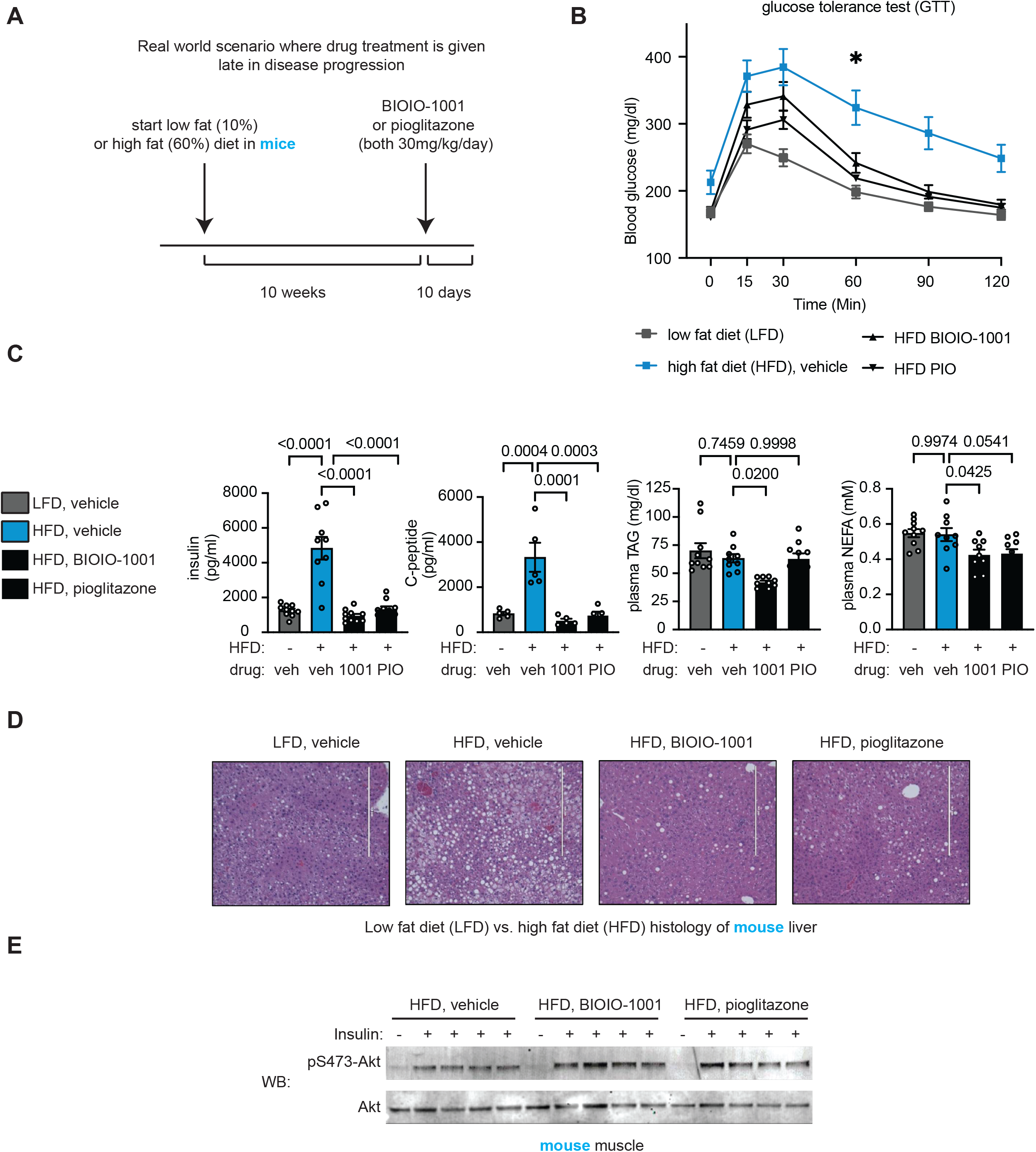
BIOIO-1001 improves systemic metabolic parameters in high fat fed mice. **A**. Schematic depicting the time course of diet and drug treatment. Mice were fed a low fat or high fat diet for 10 weeks and then treated with vehicle, Pio, or BIOIO-1001 for 10 days by gavage. **B**. Glucose tolerance testing conducted with mice fed low fat or high fat diet for 10 weeks and then treated with vehicle, Pioglitazone, or BIOIO-1001 for 8 days by gavage. **C**. Plasma insulin, c peptide, plasma triglycerides, and non-esterified fatty acids (NEFA) from animals from treated as in A. collected at sacrifice. **D**. Liver H&E histology from animals treated as in A. **E**. Western blotting was conducted with skeletal muscle lysates of mice treated as in A. and injected with a bolus of insulin or saline prior to sacrifice. Blots were probed with antibodies to total or phosphoserine 473 Akt.

## ACKNOWLEDGMENTS

We thank current and past members of the Peterson and Finck labs at WashU and at LabDAO for helpful discussions and general assistance. Especially, Damon Burrow and Nicholas Jacobs (Peterson lab) and Stanley Bishop (LabDAO).

## AUTHOR CONTRIBUTIONS

T.R.P., B.N.F., S.F., and S.Y.N. designed the study. J.L. performed the CRISPRa screen. R.F.K., S.T., and J.C. created the BIOIO-1001 series. K.G. performed nuclear receptor activation assays. K.M., N.L.B., J.P., T.P., and O.H. performed the SIRT3 genetic manipulations and accompanying in vitro experiments. K.S.M. and Y.C. performed and analyzed the HF/HTF-C diet mouse feeding experiments. J.H.H., Y.L., and J.L. performed the in vitro ALS studies. M.L.N. performed the in vivo ALS studies. T.R.P. wrote the manuscript. B.N.F., S.A.F., T.M.M., O.H., and S.Y.N. edited the manuscript.

## DATA AND MATERIALS AVAILABILITY

All data associated with this study are present in the paper or the Supplementary Materials. Shared reagents are subject to a materials transfer agreement.

## MATERIALS & METHODS

### Materials

Reagents were purchased from the following manufacturers: DMEM and RPMI 1640 Medium from Thermo Fisher Gibco, GlutaMAX Supplement from Thermo Fisher Gibco; Fetal Bovine Serum (FBS) and from Cytiva HyClone; Transit LT-1 reagent (cat. # MIR 2300) from Mirus Bio; Polyethylenimine linear MW 25000 transfection grade (PEI 25K) (cat. # 239661) from Polysciences Inc.; from Millipore Sigma; from Cayman Chemical; Bolt 4-12% Bis-Tris gels, Halt Protease Inhibitor Cocktail from Invitrogen; Bradford Reagent from Bio-Rad; from Cell Signaling Technology; Chamelon Duo Pre-Stained Ladder, IRDye 800CW and IRDye 680RD secondary antibody from Li-COR; Cell-titer Glo (cat. # G7572) from Promega; NEBNext Ultra II Q5 Master Mix (cat. # M0544L), BstXI (cat. # R0113L), Blp I (cat. # R0585L) from New England Biolabs; NucleoSpin Blood DNA isolation kit (cat. # 740950.50) from Macherey Nagel; Human Genome-wide CRISPRa-v2 Libraries were provided by Jonathan Weissman (UCSF) via Addgene (cat. # 1000000091).

BIOIO-1001 and BIOIO-1002 were synthesized by Dipharma, Inc (Kalamazoo, MI). BIOIO-1001 and BIOIO-1002 were licensed from Metabolic Solutions Development Company (MSDC, Kalamazoo, MI). UK-5099 and Pioglitazone was purchased from Sigma Chemical Co. (St. Louis, MO).

### Cell Lines and Tissue Culture

K562 CRISPRa competent cells were obtained from Johnathan Weissman. K562 cell lines were cultured in RPMI1640 Medium GlutaMAX Supplement with 10% FBS and 1% penicillin/streptomycin. Primary hepatocytes were cultured in DMEM with 10% FBS and 1% penicillin/streptomycin. G94A SOD1 iPSCs derived from PMBCs were obtained from Cedars Sinai Biomanufacturing Center (Cat. # CS2RJViALS-nxx). iPSCs were plated in Matrigel and grown in mTeSR Plus Basal Medium (Cat. # 100-0274, StemCell Technologies). All cell lines were maintained at 37°C and 5% CO_2_.

### Genome-scale screening

Genome-scale screening was carried out similar to the previously published screens ^9, 10, 41-43^. CRISPRa v2 sgRNA library ^42^ was transduced into K562 CRISPRa-competent (dCas9-SunTag) cells at a low MOI (∼0.3). Two days after transduction, the infected cells were selected with 0.75 ug/ml puromycin for three days, and the transduction was confirmed by flow cytometry. Cells were recovered from puromycin selection for two days. After two days of recovery, initial sample cells (T0; 200 million) were frozen down, and remaining cells (400 million cells) were split into either untreated (C) or treated (D) with the drug of interest. For each flask in the untreated and treated groups, the cells were kept to 0.5 million cells/ml daily. Drug treatment was continued until the untreated cells doubled five to eight more times than the treated cells. Cells were then recovered for 1 week to allow the treated cells to undergo three to four doublings. Genomic DNA isolation and library preparations were performed as previously described ^41^.

### sgRNA manipulations

sgRNA cloning for CRISPRa screen validation studies was performed according to the Weissman lab protocol: https://weissmanlab.ucsf.edu/links/sgRNACloningProtocol.pdf.

For individual validation of sgRNA phenotypes, sgRNA protospacers targeting the indicated genes or control protospacers target eGFP or non-targeting, control_1 and control_2 (we colloquially refer to these controls as “PBA392” and “PMJ051”, respectively), were individually cloned by annealing complementary synthetic oligonucleotide pairs (Integrated DNA Technologies) with flanking BstXI and BlpI restriction sites and ligating the resulting double-stranded segment into BstXI/BlpI-digested pCRISPRia-v2 (marked with a puromycin resistance cassette and BFP, Addgene #84832; ^42^). Protospacer sequences used for individual evaluation are listed below. The resulting sgRNA expression vectors were individually packaged into lentivirus. Internally controlled growth assays to evaluate sgRNA drug sensitivity phenotypes were performed by transducing cells with sgRNA expression constructs at MOI < 1 (15 – 30% infected cells), selecting to purity with puromycin (0.75μg/mL), allowing to recover for at least 1 day, treating cells with the indicated concentrations of drugs or DMSO 4-7 days after infection, and measuring the fraction of sgRNA-expressing cells 72 hours after that. During this process, populations of cells were harvested for measurement of mRNA levels by RT-qPCR (see below). These experiments were performed in triplicates from the treatment step. Knockdown of each gene was performed with their own batch of control sgRNA cells, and the data from the control cells were averaged to allow comparison of the genes on the same scale.

#### Control sgRNAs

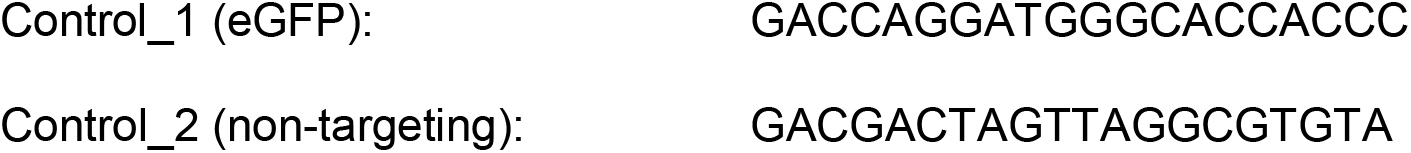

CRISPRa sgRNAs sequences below were obtained from the aforementioned CRISPRa-v2 library. We selected three out of the 10 available sgRNAs based on which ones scored best in screening.

#### Human *SIRT3*

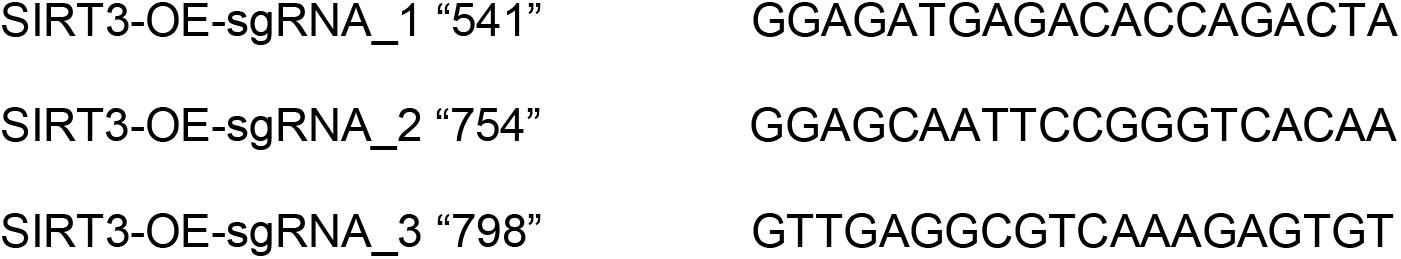

### Gene expression analysis

For individual gene analysis, total RNA was isolated, and reverse-transcription was performed from cells or tissues in the indicated conditions. Complementary DNA was made by use of a high-capacity reverse-transcription kit for liver tissues (Applied Biosystems) and a Vilo reverse-transcription kit for isolated stellate cells (Invitrogen). The resulting cDNA was diluted in Dnase-free water (1:20) followed by quantification by real-time PCR. mRNA transcripts were measured using Applied Biosystems 7900HT Sequence Detection System v2.3 software. ABI PRISM 7500 sequence detection system was also used in some cases. All data was expressed as the ratio between the expression of target gene to total RNA and/or the housekeeping genes ACTB (actin), GAPDH, or 36B4. Each treated sample was normalized to controls in the same cell/tissue type.

PCR primer sequences were obtained from: https://pga.mgh.harvard.edu/primerbank/.

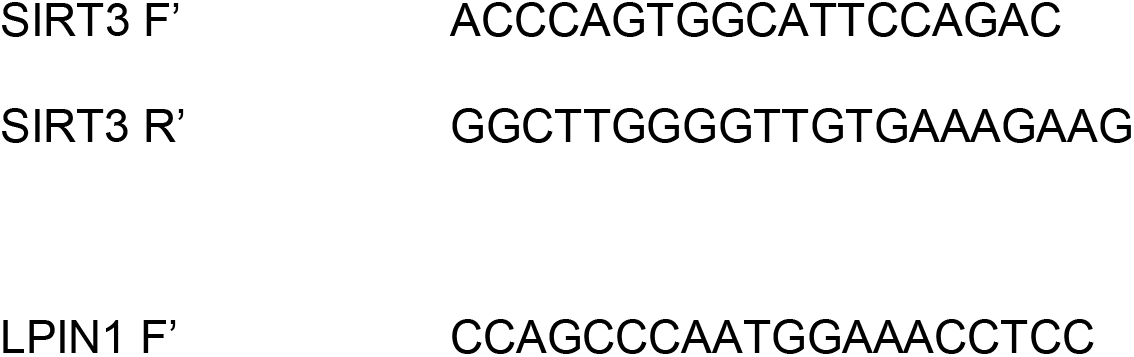
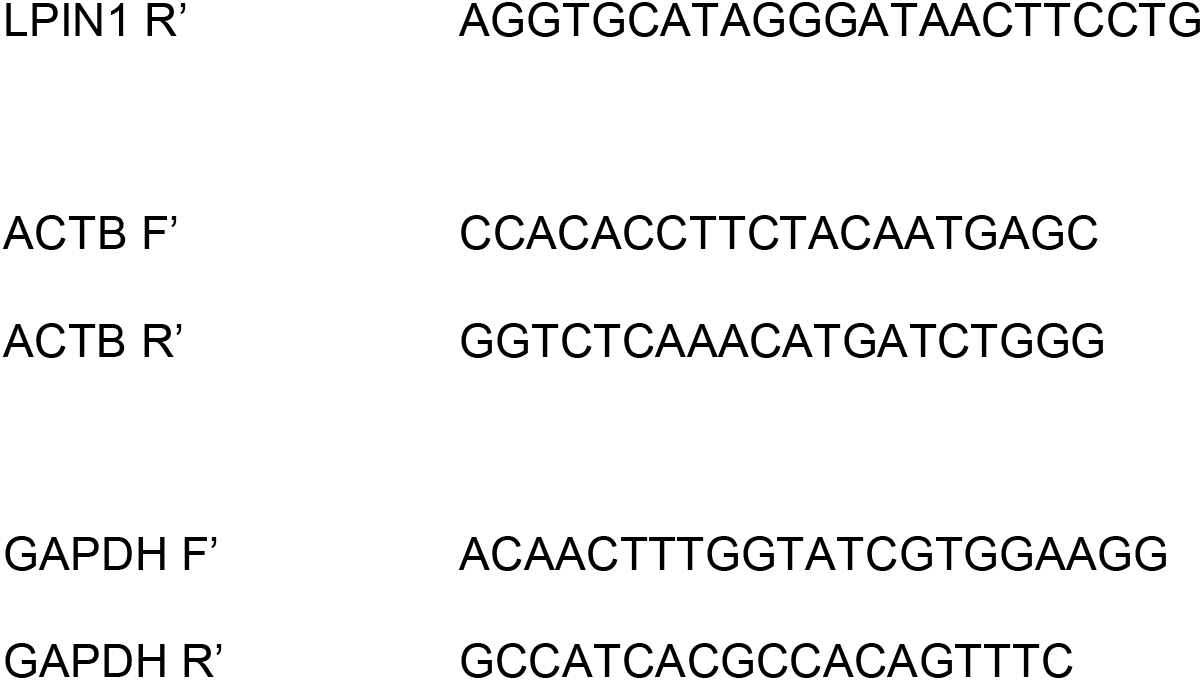

#### Mouse PCR primers

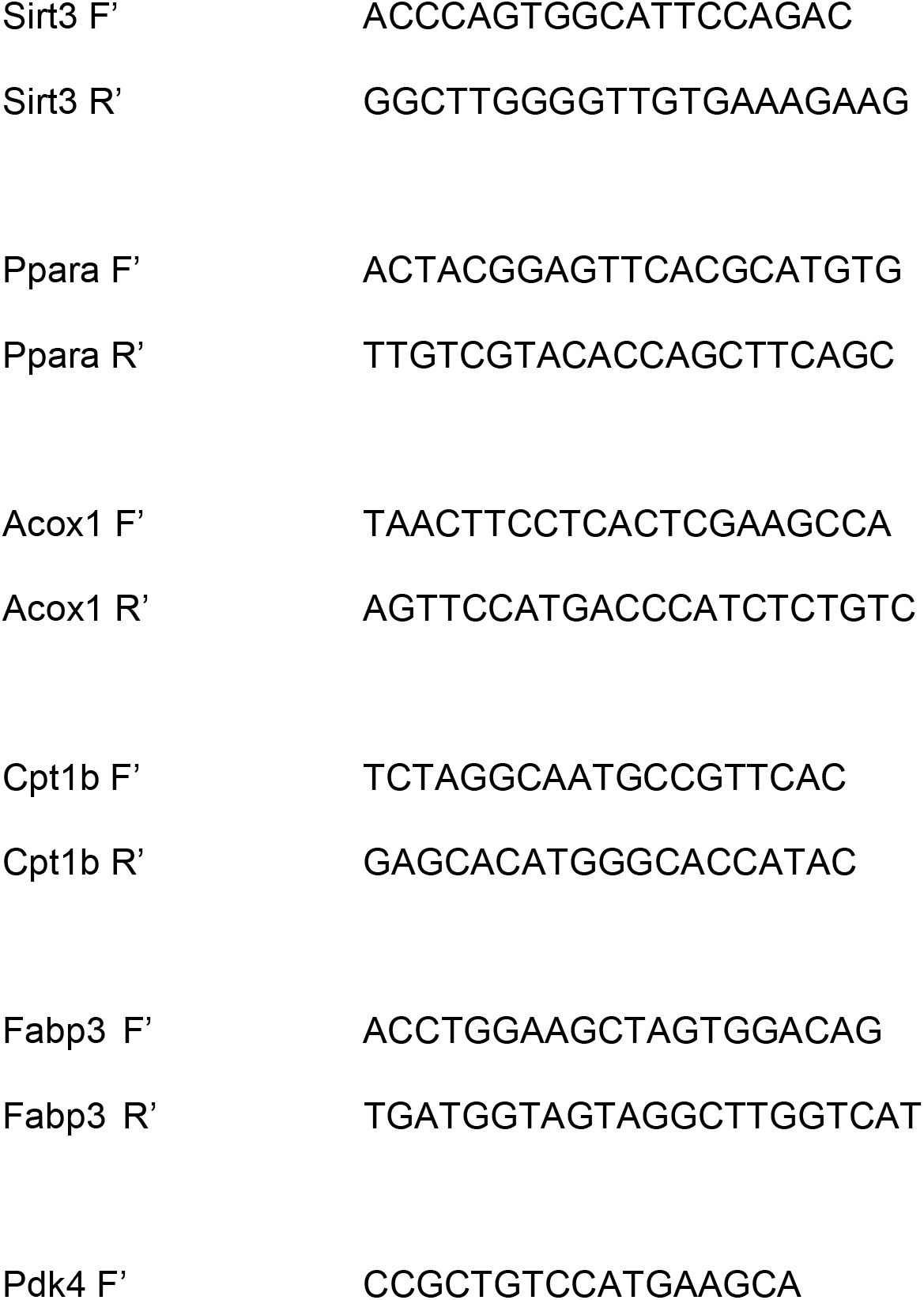
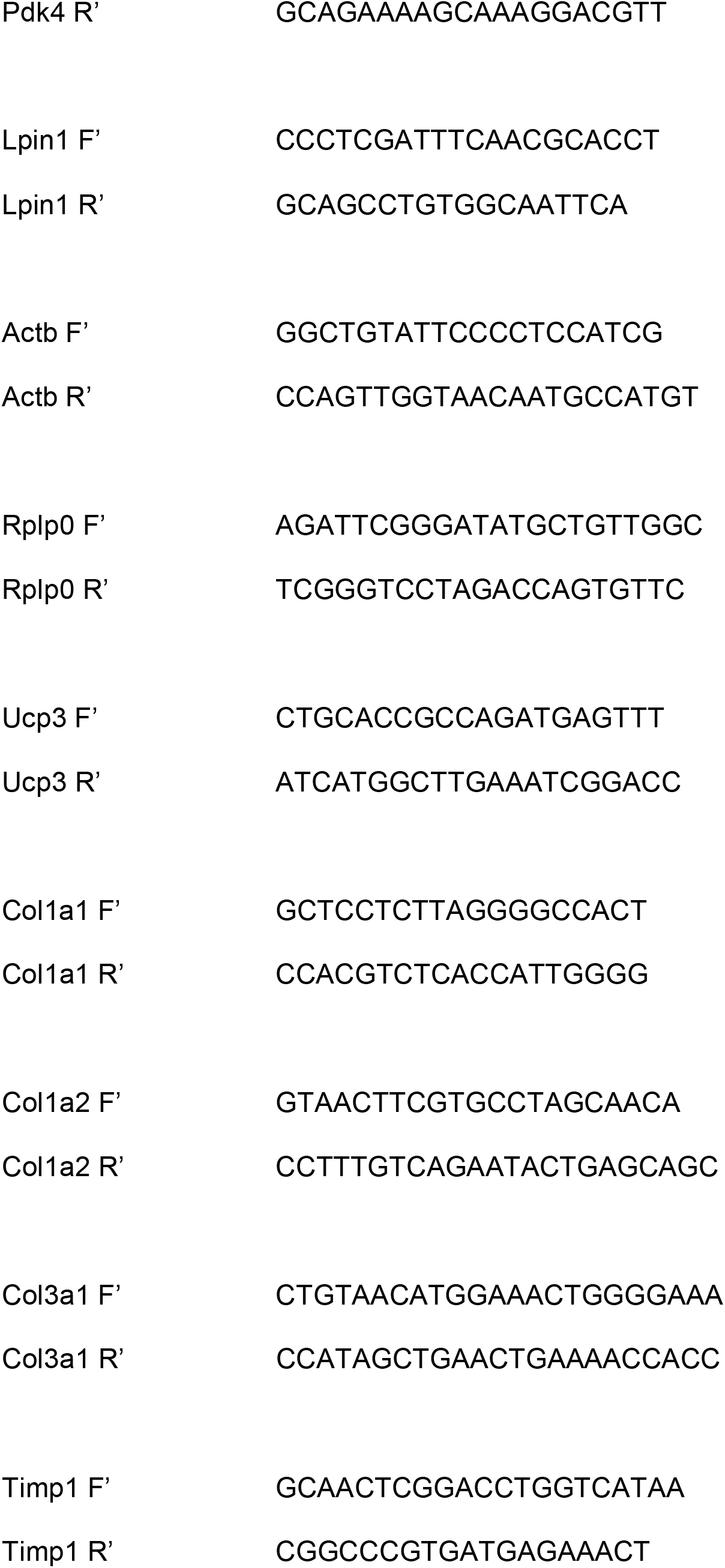
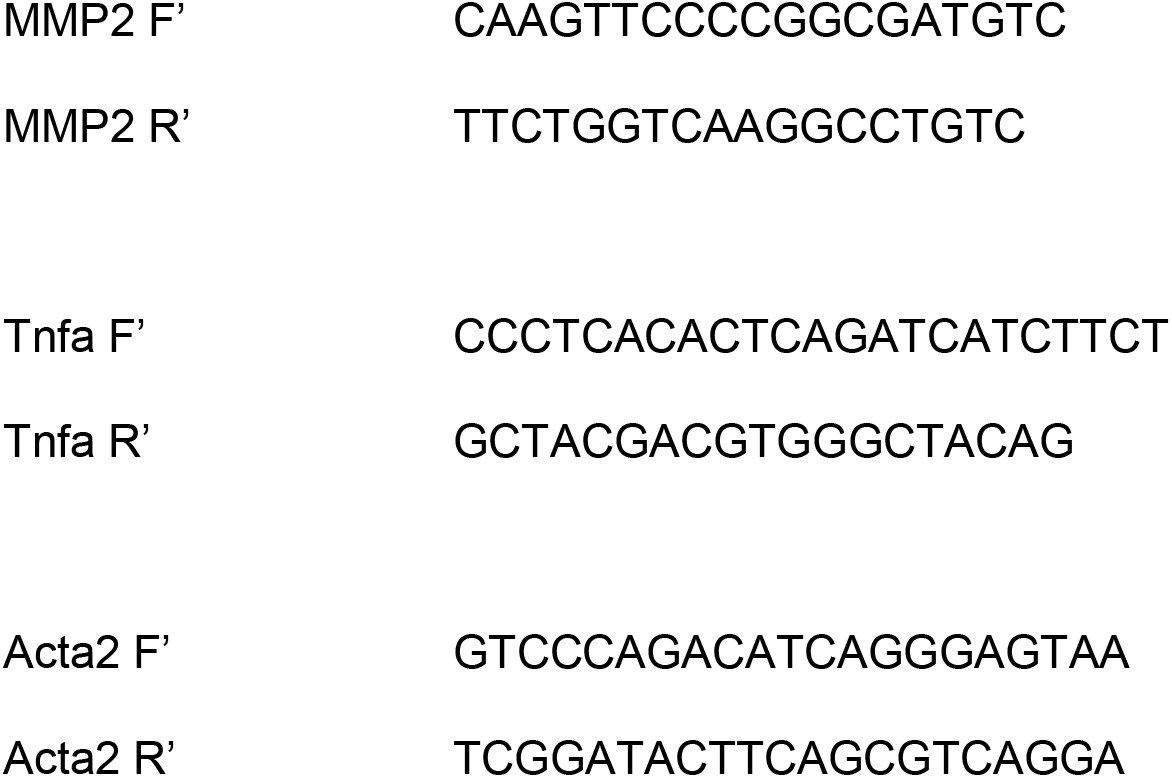

#### Genotyping primers

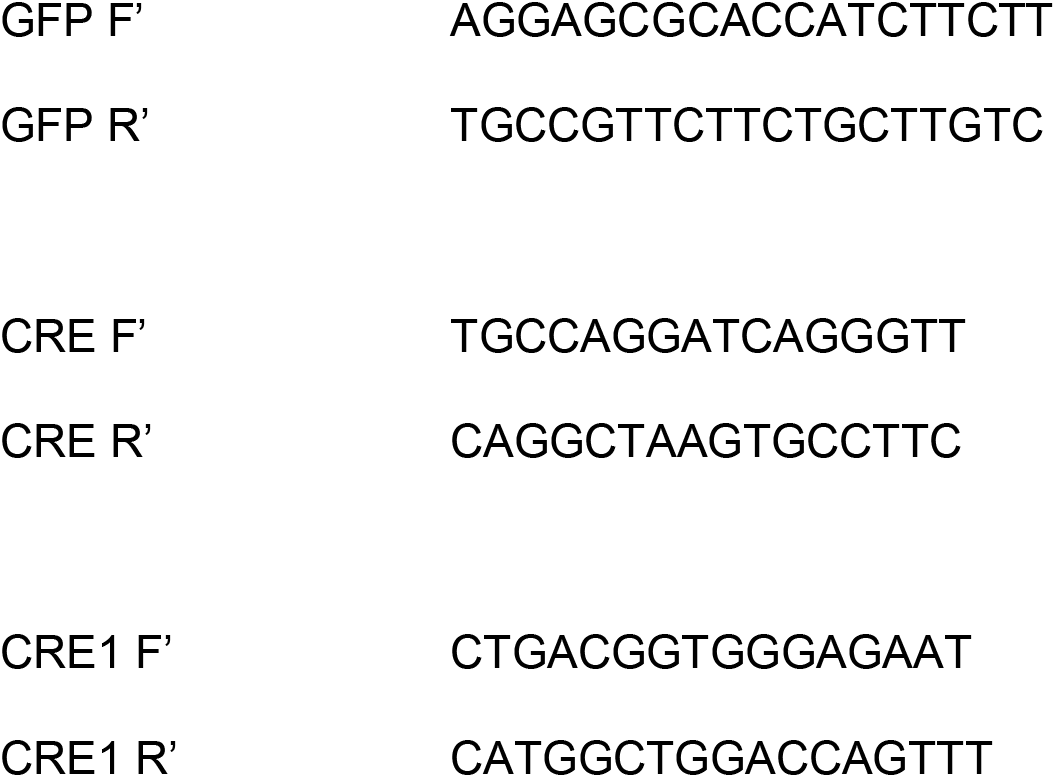

For RNA-Seq experiments, total RNA was prepared from the indicated tissues using the RNAzol method (RNA-Bee; Tel-Test) or K562 cells and analyzed using standard pipelines developed by the Genome Technology Access Center (GTAC) at Washington University.

### Cell fitness assays

Wild-type or mutant cells were seeded at 10,000 or 50,000 cells per well in a 96-well tissue culture plate and treated with indicated concentrations of compound or left untreated. 24 or 72 hours after treatment the cell viability was measured using a Cell-titer Glo colorimetric assay (Promega) according to manufacturer’s protocol. Fitness was plotted as percentage compared to untreated control. Growth Inhibitory 50% GI50 and not Inhibitory Concentration 50% IC50 was used because the latter refers to 50% inhibition of the maximal inhibition. The maximum inhibition varies for each drug, therefore using GI50 instead allowed us to compare all drugs on the same scale.

### Luciferase reporter studies

The luciferase cotransfection assays were performed in a 4-day format. HEK293 cells (ATCC) were seeded in Corning 3598 96-well plates at a density of 20,000 cells per well in 50ul of DMEM (Gibco) supplemented with 5mM L-Glutamine (Corning) and 10% FBS (Gemini Bio) and allowed to settle overnight in a 37^°^C 5% CO2 incubator. On day 2, transfection of the cells was performed by incubating Opti-MEM (Gibco), lipofectamine2000 (ThermoFisher), 100ng/ul pGL4.35[luc2P/9XGal4UAS/Hygro (Promega), and 50ng/ul chimeric Gal4-DBD fused to Nuclear Receptor-LBD in pBIND [Zeo] for 30 minutes. Twenty-five microliters of the transfection mixture was then added to the corresponding well, and the cells were gently centrifuged and placed back in the incubator overnight. The following day, cells were treated with compound or DMSO by adding 4X treatment in DMEM media with 0.4% DMSO in a volume of 25ul so that the final volume in each well was 100ul. Cells were briefly centrifuged and incubated overnight. On the final day, 75 *μ*l of OneGlo Luciferase Reagent (Promega) was added to each well and pipetted vigorously to lyse the cells. 100ul of each sample was then transferred to a Corning 3912 opaque white 96-well plate and luminescence was read on a Biotek Neo Alpha Instrument. Data was analyzed using Graphpad Prism by normalizing to DMSO (Ratio RLU Drug: DMSO) then concentrations were log-transformed, and curves were fitted by non-linear regression (agonist mode). Data is represented by mean (n = 4) +/- SEM.

### RESPYR assays

The MPC reporter system composed of human MPC2 fused to RLuc8 (BRET donor) and human MPC1 fused to Venus (BRET acceptor) was used as previously described ^44, 45^ to probe for interaction with MSDC-5514. UK-5099, a known MPC inhibitor, was used as a positive control at 5 *μ*M.

### In silico BIOIO-1001-SIRT3 docking

To generate hypotheses about the possible binding sites of BIOIO-1001 with the SIRT3 protein, diffdock ^46^, a recently developed blind docking model was used. To prepare a ligand graph, a .sdf conformer of BIOIO-1001 was generated based on BIOIO-1001’s SMILES.

A protein crystal structure measurement for SIRT3 was collected from the EBI PDBe portal. For the protein structure, we chose 4BN4 - the highest resolution structure (1.3 Angstrom) of the SIRT3 protein in complex with ADP-ribose. We removed ADP-ribose, a polyethylene glycol residue as well as Zinc and Sodium ions from the structure before blind docking.

The raw 4BN4 structure is accessible at https://www.ebi.ac.uk/pdbe/entry/pdb/4bn4 and permanently retrievable via IPFS at https://dweb.link/ipfs/bafybeihwnrfvyevmdml3ulj5wjxo6nhih5agi5t5sjwv4x7oyqyz36pgyy. The pre-processed 4BN4 structure and the BIOIO-1001 graph are retrievable at https://dweb.link/ipfs/bafybeifgdqyoydvxc2oa52cvfsogk4l6zblybvqejeatstcgg3lrbj6ocu and https://dweb.link/ipfs/bafybeie4d6kpkk47c6njn6nko4jxo4i5aaie46gxacph7lituqitf2q2ca, respectively.

We ran diffdock with default parameters and visualized the top 10 ligand poses with color-coded confidence scores. An interactive version of diffdock can be accessed at https://colab.research.google.com/drive/1nvCyQkbO-TwXZKJ0RCShVEym1aFWxlkX.

The docked results are retrievable via: https://dweb.link/ipfs/bafybeibidrz2vef7uc7hslh2tauuu2rgwjx7dld6dng3zrfodla6uyyq6e.

### Animal studies

All animal experiments were approved by the Institutional Animal Care and Use Committees of Washington University and Saint Louis University and comply with the criteria outlined in the Guide for the Care and Use of Laboratory Animals by the National Academy of Sciences. All diet studies were initiated in mice 6-9 weeks old. Wild-type C57BL6/J and SV129 control mice and SIRT3 and G93A SOD1 mutant mice were purchased from Jackson Laboratories.

To induce diet induced obesity, six- to 8-week-old mice were placed on purified diet providing 10% (Research Diets Inc., D09100304) or 60% (Research Diets Inc., D12492) of its calories as fat and remained on these diets for 10 weeks before initiation of drug administration. Mice were then treated with BIOIO-1001 (30 mg/kg/day), PIO (30 mg/kg/day), or vehicle control by daily oral gavage for 1 week prior to euthanasia. Vehicle gavage solution was composed of 1% low-viscosity carboxymethylcellulose, 0.1% Tween-80, and 5% DMSO.

For studies evaluating the effects of BIOIO-1001 or Pio on liver injury and NASH endpoints, male mice were placed on a diet enriched with fat (40% kcal, mainly trans-fat; trans-oleic and trans-linoleic acids), fructose (20% kcal), and cholesterol (2% w/w) (HTF-C diet; D17010103); Research Diets Inc.). Control mice were fed a matched low-fat (LF; 10% kcal) diet that was not supplemented with fructose or cholesterol (D12450J; Research Diets Inc.). Mice were treated with vehicle or BIOIO-1001 by daily oral gavage for 3 weeks prior to sacrifice.

Glucose tolerance tests (GTT) were conducted in mice after 8 days of treatment with vehicle, Pio, or BIOIO-1001. Mice were fasted overnight for 16 h on aspen chip bedding and GTTs were performed as previously reported ^47^. Blood glucose area under the curve (AUC) was calculated using the trapezoidal rule.

Mice were sacrificed by CO_2_ asphyxiation and tissues and plasma were collected after a 4-hour fast. Liver samples were weighed, frozen in liquid nitrogen, and stored at −80°C. To examine insulin-stimulated insulin signaling, mice were fasted overnight and injected intravenously with human insulin (10mU/g body weight) as described ^47^ 5 minutes prior to sacrifice.

### Plasma chemistry

Insulin content was analyzed by Singulex assay by the Washington University Immunoassay Core of the Diabetes Research Center. Plasma triglyceride and cholesterol concentrations were measured by Infinity colorimetric assay kits (Thermo Fisher Scientific). Plasma non-esterified fatty acids concentrations were measured using enzymatic assay (Wako Diagnostics). Adiponectin concentrations were determined by using ELISA (Millipore). Plasma alanine transaminase (ALT) and aspartate transaminase (AST) were measured by kinetic absorbance assays (Teco Diagnostics).

### Protein isolation and western blotting analyses

Protein from frozen tissue was homogenized in 0.3-1 ml ice-cold homogenization buffer (25mM HEPES, 150 mM NaCl, 5 mM EDTA, 1% Triton X-100, pH 8.0, supplemented with 1 mM activated Na_3_VO_4_, 1 mM phenylmethanesulfonyl fluoride, 5 mM sodium fluoride, and 1X Complete protease inhibitor cocktail tablet (Roche, Manneheim, Germany; cat. 04693116001) using high-speed tissue disruption with the TissueLyser II (Qiagen, Valencia, CA). Tissue homogenates were subsequently solubilized by rotating at 4°C at 50 rpm for 1 h before being centrifuged (15,000 g for 15 min at 4°C) and collecting the supernatant.

Lysates were subjected to SDS-PAGE and transferred to PVDF membranes. Blots were then rinsed with Tris-buffered saline plus Tween (TBST) (0.14 mol/l NaCl, 0.02 mol/l Tris base, pH 7.6, and 0.1% Tween), blocked with 5% BSA in TBST for 1 h at room temperature, washed 3 × 10 min at room temperature, and incubated with the relevant primary antibody 1:1000 in 5% BSA overnight at 4°C. Blots were then washed 3 × 5 min with TBST, incubated with relevant secondary antibodies for 1 h at room temperature, washed again 3 × 10 min with TBST, and washed 2 × 10 min with TBS. Protein bands were visualized using the Odyssey Imaging System (LiCor Biosciences, Lincoln, NE). Akt (cat. 9272) and phospho-Akt Ser473 (cat. 9271) antibodies were obtained from Cell Signaling (Danvers, MA). Goat anti-rabbit 800 (cat. 926-32211) secondary antibodies were obtained from LiCor Biosciences (Lincoln, NE).

### Histologic scoring

Liver sections were fixed in 10% neutral buffered formalin for 24 hours and then embedded in paraffin blocks. Sections were cut and stained with either hematoxylin and eosin or Masson’s trichrome stain. Livers were analyzed by a liver pathologist, blinded to treatment group and genotype. Steatosis, inflammation, hepatocyte ballooning, and fibrosis were scored according to NAFLD activity score (NAS) and fibrosis scoring ^48^.

### Motor neuron differentiation from human induced pluripotent stem cells

Human induced pluripotent stem cell lines were routinely cultured on Matrigel-coated dishes in MACS iPS-Brew media (Miltenyi Biotec) and passaged using ReLESR (Stem Cell Technologies) on a weekly basis. To induce motor neuron differentiation, pluripotent colonies were detached from the culture dish using Accutase and exposed to neural induction media consisting of 4.25 μM CHIR99021 and 0.5 μM LDN-193189 for the first 10 days for culture on Matrigel-coated dishes. 1 μM Retinoic acid (RA) was supplemented into this media from days 3 to 10. From days 10 to 17, cells were cultured in motor neuron patterning media consisting of 1 μM RA and 1 μM Purmorphamine. Subsequently, from days 18 to 28, the adherent culture was dissociated into single cells and re-plated onto Matrigel-coated dishes in media consisting of 10 ng/ml BDNF and 10 ng/ml GDNF. N2B27 media: 50% DMEM/F12, 50% Neurobasal medium, 1X N2 supplement, 1X B27 supplement, 1X NEAA and 1X Glutamax.

### BIOIO-1001 treatment in sporadic ALS motor neuron cultures

Sporadic ALS motor neuron cultures were dissociated with Accutase and seeded at 100,000 cells per well in a 96-well plate on day 27. On day 28, cells were treated with BIOIO-1001 at the desired concentrations of 1 μM,5 μM and 10 μM. Cells were then fixed for immunostaining 3 days after BIO1O-1001 treatment.

### Immunofluorescence, image acquisition and image analysis of iPSC-derived motor neurons

Cells were fixed in 4% paraformaldehyde for 15 min, permeabilized in 0.1% Triton X-100 for 15 minutes and blocked in buffer containing 5% FBS and 1% BSA for an hour at room temperature. Primary antibodies were diluted in blocking buffer and incubated overnight at 4 °C. The following antibodies (and their respective dilutions) were used: rabbit ISL1 (Abcam ab109517; 1:1000), mouse SMI-32 (BioLegend 801701; 1:1000). The respective secondary antibodies (Molecular Probes, Invitrogen) were diluted at 1:1500 in blocking buffer and incubated at room temperature, in the dark, for 90 minutes. DAPI (0.1μg/ml) to visualize cellular nuclei. Images were acquired using the high content microscope Opera Phenix (Perkin Elmer) using the 20x air objective. Image analyses including cell counts and intensity measurements were performed using Columbus (Perkin Elmer).

### ALS phenotype scoring

The onset of ALS disease was determined by peak body weight in conjunction with neurological scoring. Each week body weights were recorded, and neurological scoring was performed. Score criteria range from 0 (good neurological function) to 4 (poor neurological function). The difference between age of onset and euthanasia was used as a measurement of disease progression.

### Statistical Analyses

P values for comparing more than two groups were calculated using ANOVA coupled to Tukey’s multiple comparison tests. *P* values for RESPYR curves were calculated using repeated measures, ANOVA coupled to Tukey’s multiple comparison tests. *P* values for pairwise comparisons were calculated using a Student’s t-test. In all experiments, *P* ≤ 0.05 was used to determine significant difference. All quantitative data is represented as mean ± SEM.

## REFERENCES

1. Zheng W, Thorne N, McKew JC. Phenotypic screens as a renewed approach for drug discovery. Drug Discov Today. 2013;18(21-22):1067–73. Epub 2013/07/16. doi: 10.1016/j.drudis.2013.07.001. PubMed PMID: 23850704; PMCID: PMC4531371.

2. Jost M, Weissman JS. CRISPR Approaches to Small Molecule Target Identification. ACS Chem Biol. 2018;13(2):366–75. doi: 10.1021/acschembio.7b00965. PubMed PMID: 29261286; PMCID: PMC5834945.

3. Lopez-Otin C, Blasco MA, Partridge L, Serrano M, Kroemer G. The hallmarks of aging. Cell. 2013;153(6):1194–217. doi: 10.1016/j.cell.2013.05.039. PubMed PMID: 23746838; PMCID: PMC3836174.

4. Finkel T. The metabolic regulation of aging. Nat Med. 2015;21(12):1416–23. Epub 2015/12/10. doi: 10.1038/nm.3998. PubMed PMID: 26646498.

5. Barzilai N, Ferrucci L. Insulin resistance and aging: a cause or a protective response? J Gerontol A Biol Sci Med Sci. 2012;67(12):1329–31. Epub 2012/08/04. doi: 10.1093/gerona/gls145. PubMed PMID: 22859390.

6. Davidson MA, Mattison DR, Azoulay L, Krewski D. Thiazolidinedione drugs in the treatment of type 2 diabetes mellitus: past, present and future. Crit Rev Toxicol. 2018;48(1):52–108. Epub 2017/08/18. doi: 10.1080/10408444.2017.1351420. PubMed PMID: 28816105.

7. Lehmann JM, Moore LB, Smith-Oliver TA, Wilkison WO, Willson TM, Kliewer SA. An antidiabetic thiazolidinedione is a high affinity ligand for peroxisome proliferator-activated receptor gamma (PPAR gamma). J Biol Chem. 1995;270(22):12953–6. doi: 10.1074/jbc.270.22.12953. PubMed PMID: 7768881.

8. Colca JR, McDonald WG, Cavey GS, Cole SL, Holewa DD, Brightwell-Conrad AS, Wolfe CL, Wheeler JS, Coulter KR, Kilkuskie PM, Gracheva E, Korshunova Y, Trusgnich M, Karr R, Wiley SE, Divakaruni AS, Murphy AN, Vigueira PA, Finck BN, Kletzien RF. Identification of a mitochondrial target of thiazolidinedione insulin sensitizers (mTOT)--relationship to newly identified mitochondrial pyruvate carrier proteins. PLoS One. 2013;8(5):e61551. Epub 20130515. doi: 10.1371/journal.pone.0061551. PubMed PMID: 23690925; PMCID: PMC3655167.

9. Jost M, Chen Y, Gilbert LA, Horlbeck MA, Krenning L, Menchon G, Rai A, Cho MY, Stern JJ, Prota AE, Kampmann M, Akhmanova A, Steinmetz MO, Tanenbaum ME, Weissman JS. Combined CRISPRi/a-Based Chemical Genetic Screens Reveal that Rigosertib Is a Microtubule-Destabilizing Agent. Mol Cell. 2017;68(1):210–23 e6. doi: 10.1016/j.molcel.2017.09.012. PubMed PMID: 28985505; PMCID: PMC5640507.

10. Yu Z, Surface LE, Park CY, Horlbeck MA, Wyant GA, Abu-Remaileh M, Peterson TR, Sabatini DM, Weissman JS, O’Shea EK. Identification of a transporter complex responsible for the cytosolic entry of nitrogen-containing-bisphosphonates. Elife. 2018;7. doi: 10.7554/eLife.36620. PubMed PMID: 29745899.

11. Kumar S, Li J, Park J, Hart SK, Song NJ, Burrow DT, Bean NL, Jacobs NC, Coler-Reilly A, Pendergrass AO, Pierre TH, Bradley IC, Carette JE, Varadarajan M, Brummelkamp TR, Dolle R, Peterson TR. Sphingolipid Biosynthesis Inhibition As A Host Strategy Against Diverse Pathogens. bioRxiv. 2020:2020.04.10.035683. doi: 10.1101/2020.04.10.035683.

12. Finck BN, Gropler MC, Chen Z, Leone TC, Croce MA, Harris TE, Lawrence JC, Jr., Kelly DP. Lipin 1 is an inducible amplifier of the hepatic PGC-1alpha/PPARalpha regulatory pathway. Cell Metab. 2006;4(3):199–210. doi: 10.1016/j.cmet.2006.08.005. PubMed PMID: 16950137.

13. Hirschey MD, Shimazu T, Goetzman E, Jing E, Schwer B, Lombard DB, Grueter CA, Harris C, Biddinger S, Ilkayeva OR, Stevens RD, Li Y, Saha AK, Ruderman NB, Bain JR, Newgard CB, Farese RV, Jr., Alt FW, Kahn CR, Verdin E. SIRT3 regulates mitochondrial fatty-acid oxidation by reversible enzyme deacetylation. Nature. 2010;464(7285):121–5. doi: 10.1038/nature08778. PubMed PMID: 20203611; PMCID: PMC2841477.

14. Caron A, Richard D, Laplante M. The Roles of mTOR Complexes in Lipid Metabolism. Annu Rev Nutr. 2015;35:321–48. Epub 2015/07/18. doi: 10.1146/annurev-nutr-071714-034355. PubMed PMID: 26185979.

15. Katsyuba E, Romani M, Hofer D, Auwerx J. NAD(+) homeostasis in health and disease. Nat Metab. 2020;2(1):9–31. Epub 2020/07/23. doi: 10.1038/s42255-019-0161-5. PubMed PMID: 32694684.

16. Dwyer JR, Donkor J, Zhang P, Csaki LS, Vergnes L, Lee JM, Dewald J, Brindley DN, Atti E, Tetradis S, Yoshinaga Y, De Jong PJ, Fong LG, Young SG, Reue K. Mouse lipin-1 and lipin-2 cooperate to maintain glycerolipid homeostasis in liver and aging cerebellum. Proc Natl Acad Sci U S A. 2012;109(37):E2486–95. Epub 20120820. doi: 10.1073/pnas.1205221109. PubMed PMID: 22908270; PMCID: PMC3443145.

17. Kuleshov MV, Jones MR, Rouillard AD, Fernandez NF, Duan Q, Wang Z, Koplev S, Jenkins SL, Jagodnik KM, Lachmann A, McDermott MG, Monteiro CD, Gundersen GW, Ma’ayan A. Enrichr: a comprehensive gene set enrichment analysis web server 2016 update. Nucleic Acids Res. 2016;44(W1):W90–7. Epub 2016/05/05. doi: 10.1093/nar/gkw377. PubMed PMID: 27141961; PMCID: PMC4987924.

18. Akinwumi BC, Bordun KM, Anderson HD. Biological Activities of Stilbenoids. Int J Mol Sci. 2018;19(3). Epub 2018/03/10. doi: 10.3390/ijms19030792. PubMed PMID: 29522491; PMCID: PMC5877653.

19. Howitz KT, Bitterman KJ, Cohen HY, Lamming DW, Lavu S, Wood JG, Zipkin RE, Chung P, Kisielewski A, Zhang LL, Scherer B, Sinclair DA. Small molecule activators of sirtuins extend Saccharomyces cerevisiae lifespan. Nature. 2003;425(6954):191–6. Epub 20030824. doi: 10.1038/nature01960. PubMed PMID: 12939617.

20. Saxton RA, Sabatini DM. mTOR Signaling in Growth, Metabolism, and Disease. Cell. 2017;169(2):361–71. doi: 10.1016/j.cell.2017.03.035. PubMed PMID: 28388417.

21. Covarrubias AJ, Perrone R, Grozio A, Verdin E. NAD(+) metabolism and its roles in cellular processes during ageing. Nat Rev Mol Cell Biol. 2021;22(2):119–41. Epub 2020/12/24. doi: 10.1038/s41580-020-00313-x. PubMed PMID: 33353981; PMCID: PMC7963035.

22. Schwabe RF, Tabas I, Pajvani UB. Mechanisms of Fibrosis Development in Nonalcoholic Steatohepatitis. Gastroenterology. 2020;158(7):1913–28. Epub 2020/02/12. doi: 10.1053/j.gastro.2019.11.311. PubMed PMID: 32044315; PMCID: PMC7682538.

23. Schuster S, Cabrera D, Arrese M, Feldstein AE. Triggering and resolution of inflammation in NASH. Nat Rev Gastroenterol Hepatol. 2018;15(6):349–64. Epub 2018/05/10. doi: 10.1038/s41575-018-0009-6. PubMed PMID: 29740166.

24. Tryka KA, Hao L, Sturcke A, Jin Y, Wang ZY, Ziyabari L, Lee M, Popova N, Sharopova N, Kimura M, Feolo M. NCBI’s Database of Genotypes and Phenotypes: dbGaP. Nucleic Acids Res. 2014;42(Database issue):D975–9. Epub 2013/12/04. doi: 10.1093/nar/gkt1211. PubMed PMID: 24297256; PMCID: PMC3965052.

25. Li Z, Shi L, Li X, Wang X, Wang H, Liu Y. RNF144A-AS1, a TGF-beta1- and hypoxia-inducible gene that promotes tumor metastasis and proliferation via targeting the miR-30c-2-3p/LOX axis in gastric cancer. Cell Biosci. 2021;11(1):177. Epub 20210928. doi: 10.1186/s13578-021-00689-z. PubMed PMID: 34583752; PMCID: PMC8480077.

26. Yang L, Han B, Zhang M, Wang YH, Tao K, Zhu MX, He K, Zhang ZG, Hou S. Activation of BK Channels Prevents Hepatic Stellate Cell Activation and Liver Fibrosis Through the Suppression of TGFbeta1/SMAD3 and JAK/STAT3 Profibrotic Signaling Pathways. Front Pharmacol. 2020;11:165. Epub 20200306. doi: 10.3389/fphar.2020.00165. PubMed PMID: 32210801; PMCID: PMC7068464.

27. Hall RA, Liebe R, Hochrath K, Kazakov A, Alberts R, Laufs U, Bohm M, Fischer HP, Williams RW, Schughart K, Weber SN, Lammert F. Systems genetics of liver fibrosis: identification of fibrogenic and expression quantitative trait loci in the BXD murine reference population. PLoS One. 2014;9(2):e89279. Epub 20140228. doi: 10.1371/journal.pone.0089279. PubMed PMID: 24586654; PMCID: PMC3938463.

28. Wang P, Dai X, Jiang W, Li Y, Wei W. RBR E3 ubiquitin ligases in tumorigenesis. Semin Cancer Biol. 2020;67(Pt 2):131–44. Epub 20200519. doi: 10.1016/j.semcancer.2020.05.002. PubMed PMID: 32442483.

29. Apolloni S, D’Ambrosi N. Fibrosis as a common trait in amyotrophic lateral sclerosis tissues. Neural Regen Res. 2022;17(1):97–8. Epub 2021/06/09. doi: 10.4103/1673-5374.314302. PubMed PMID: 34100438; PMCID: PMC8451558.

30. Steyn FJ, Li R, Kirk SE, Tefera TW, Xie TY, Tracey TJ, Kelk D, Wimberger E, Garton FC, Roberts L, Chapman SE, Coombes JS, Leevy WM, Ferri A, Valle C, René F, Loeffler JP, McCombe PA, Henderson RD, Ngo ST. Altered skeletal muscle glucose-fatty acid flux in amyotrophic lateral sclerosis. Brain Commun. 2020;2(2):fcaa154. Epub 2020/11/27. doi: 10.1093/braincomms/fcaa154. PubMed PMID: 33241210; PMCID: PMC7677608.

31. Masrori P, Van Damme P. Amyotrophic lateral sclerosis: a clinical review. Eur J Neurol. 2020;27(10):1918–29. Epub 2020/06/12. doi: 10.1111/ene.14393. PubMed PMID: 32526057; PMCID: PMC7540334.

32. Gurney ME, Pu H, Chiu AY, Dal Canto MC, Polchow CY, Alexander DD, Caliendo J, Hentati A, Kwon YW, Deng HX, et al. Motor neuron degeneration in mice that express a human Cu,Zn superoxide dismutase mutation. Science. 1994;264(5166):1772–5. Epub 1994/06/17. doi: 10.1126/science.8209258. PubMed PMID: 8209258.

33. Dupuis L, Corcia P, Fergani A, Gonzalez De Aguilar JL, Bonnefont-Rousselot D, Bittar R, Seilhean D, Hauw JJ, Lacomblez L, Loeffler JP, Meininger V. Dyslipidemia is a protective factor in amyotrophic lateral sclerosis. Neurology. 2008;70(13):1004–9. Epub 20080116. doi: 10.1212/01.wnl.0000285080.70324.27. PubMed PMID: 18199832.

34. Hornsby PJ. Chapter 4 - The nature of aging and the geroscience hypothesis. In: Musi N, Hornsby PJ, editors. Handbook of the Biology of Aging (Ninth Edition): Academic Press; 2021. p. 69–76.

35. Neganova ME, Klochkov SG, Aleksandrova YR, Aliev G. The Hydroxamic Acids as Potential Anticancer and Neuroprotective Agents. Curr Med Chem. 2021;28(39):8139–62. doi: 10.2174/0929867328666201218123154. PubMed PMID: 33342403.

36. Li TY, Song L, Sun Y, Li J, Yi C, Lam SM, Xu D, Zhou L, Li X, Yang Y, Zhang CS, Xie C, Huang X, Shui G, Lin SY, Reue K, Lin SC. Tip60-mediated lipin 1 acetylation and ER translocation determine triacylglycerol synthesis rate. Nat Commun. 2018;9(1):1916. Epub 20180515. doi: 10.1038/s41467-018-04363-w. PubMed PMID: 29765047; PMCID: PMC5953937.

37. Jing H, Lin H. Sirtuins in epigenetic regulation. Chem Rev. 2015;115(6):2350–75. Epub 20150128. doi: 10.1021/cr500457h. PubMed PMID: 25804908; PMCID: PMC4610301.

38. Xie N, Zhang L, Gao W, Huang C, Huber PE, Zhou X, Li C, Shen G, Zou B. NAD(+) metabolism: pathophysiologic mechanisms and therapeutic potential. Signal Transduct Target Ther. 2020;5(1):227. Epub 20201007. doi: 10.1038/s41392-020-00311-7. PubMed PMID: 33028824; PMCID: PMC7539288.

39. Woodcock HV, Eley JD, Guillotin D, Plate M, Nanthakumar CB, Martufi M, Peace S, Joberty G, Poeckel D, Good RB, Taylor AR, Zinn N, Redding M, Forty EJ, Hynds RE, Swanton C, Karsdal M, Maher TM, Fisher A, Bergamini G, Marshall RP, Blanchard AD, Mercer PF, Chambers RC. The mTORC1/4E-BP1 axis represents a critical signaling node during fibrogenesis. Nat Commun. 2019;10(1):6. Epub 20190102. doi: 10.1038/s41467-018-07858-8. PubMed PMID: 30602778; PMCID: PMC6315032.

40. Mitchell SJ, Scheibye-Knudsen M, Longo DL, de Cabo R. Animal models of aging research: implications for human aging and age-related diseases. Annu Rev Anim Biosci. 2015;3:283–303. Epub 2015/02/18. doi: 10.1146/annurev-animal-022114-110829. PubMed PMID: 25689319.

41. Gilbert LA, Horlbeck MA, Adamson B, Villalta JE, Chen Y, Whitehead EH, Guimaraes C, Panning B, Ploegh HL, Bassik MC, Qi LS, Kampmann M, Weissman JS. Genome-Scale CRISPR-Mediated Control of Gene Repression and Activation. Cell. 2014;159(3):647–61. doi: 10.1016/j.cell.2014.09.029. PubMed PMID: 25307932; PMCID: PMC4253859.

42. Horlbeck MA, Gilbert LA, Villalta JE, Adamson B, Pak RA, Chen Y, Fields AP, Park CY, Corn JE, Kampmann M, Weissman JS. Compact and highly active next-generation libraries for CRISPR-mediated gene repression and activation. Elife. 2016;5. doi: 10.7554/eLife.19760. PubMed PMID: 27661255; PMCID: PMC5094855.

43. Horlbeck MA, Xu A, Wang M, Bennett NK, Park CY, Bogdanoff D, Adamson B, Chow ED, Kampmann M, Peterson TR, Nakamura K, Fischbach MA, Weissman JS, Gilbert LA. Mapping the Genetic Landscape of Human Cells. Cell. 2018;174(4):953–67 e22. Epub 2018/07/24. doi: 10.1016/j.cell.2018.06.010. PubMed PMID: 30033366; PMCID: PMC6426455.

44. Vigueira PA, McCommis KS, Hodges WT, Schweitzer GG, Cole SL, Oonthonpan L, Taylor EB, McDonald WG, Kletzien RF, Colca JR, Finck BN. The beneficial metabolic effects of insulin sensitizers are not attenuated by mitochondrial pyruvate carrier 2 hypomorphism. Exp Physiol. 2017;102(8):985–99. doi: 10.1113/EP086380. PubMed PMID: 28597936; PMCID: PMC5667918.

45. McCommis KS, Hodges WT, Brunt EM, Nalbantoglu I, McDonald WG, Holley C, Fujiwara H, Schaffer JE, Colca JR, Finck BN. Targeting the mitochondrial pyruvate carrier attenuates fibrosis in a mouse model of nonalcoholic steatohepatitis. Hepatology. 2017;65(5):1543–56. doi: 10.1002/hep.29025. PubMed PMID: 28027586; PMCID: PMC5397348.

46. Gabriele Corso HS, Bowen Jing, Regina Barzilay, Tommi Jaakkola. DiffDock: Diffusion Steps, Twists, and Turns for Molecular Docking. Available from: https://arxiv.org/abs/2210.01776.

47. Finck BN, Bernal-Mizrachi C, Han DH, Coleman T, Sambandam N, LaRiviere LL, Holloszy JO, Semenkovich CF, Kelly DP. A potential link between muscle peroxisome proliferator-activated receptor-alpha signaling and obesity-related diabetes. Cell Metab. 2005;1(2):133–44. Epub 2005/08/02. doi: 10.1016/j.cmet.2005.01.006. PubMed PMID: 16054054.

48. Kleiner DE, Brunt EM, Van Natta M, Behling C, Contos MJ, Cummings OW, Ferrell LD, Liu YC, Torbenson MS, Unalp-Arida A, Yeh M, McCullough AJ, Sanyal AJ, Nonalcoholic Steatohepatitis Clinical Research N. Design and validation of a histological scoring system for nonalcoholic fatty liver disease. Hepatology. 2005;41(6):1313–21. doi: 10.1002/hep.20701. PubMed PMID: 15915461.

